# Passenger DNA alterations reduce cancer fitness in cell culture and mouse models

**DOI:** 10.1101/026302

**Authors:** Christopher D McFarland, Julia A Yaglom, Jonathan W Wojtkowiak, Jacob G Scott, David L Morse, Michael Y Sherman, Leonid A Mirny

## Abstract

Genomic instability causes cancers to acquire hundreds to thousands of mutations and chromosomal alterations during their somatic evolution. Most of these mutations and alterations are termed passengers because they do not confer cancer phenotypes. Evolutionary simulations and cancer genomic studies suggested that mildly-deleterious passengers accumulate, collectively slow cancer progression, reduce the fitness of cancer cells and enhance the effects of therapeutics. However, these effects of passengers and their impact on clinical variables remain limited to genomic analysis. Here, to assess passengers’ effect on cell fitness and cancer, we specifically introduced increasing passenger loads into human cell lines and mouse models. We found that passenger load dramatically reduced cancer cell’s fitness in every model investigated. Passengers’ average fitness cost of ∼3% per MB, indicates that genomic instability in cancer in patients can slow tumor growth and prevent metastatic progression. We conclude that genomic instability in cancer is a double-edged sword: it accelerates the accumulation of adaptive drivers, yet incurs a harmful passenger load that can outweigh drivers’ benefit. Passenger load could be a useful biomarker for tumor aggressiveness and response to mutagenic or passenger-exacerbating therapies, including anti-tumor immunity.

## Introduction

Genomic instability, i.e. a high frequency of mutations and chromosomal alterations (referred to collectively as mutations) within cellular lineages, is a hallmark of carcinogenesis^1^. Genomic instability can create *driver mutations* that confer a fitness advantage to somatic cells in their microenvironment and expand (i.e. *drive*) the cell lineage to cancer. However, drivers are rare. Indeed, 97% of all somatic cancer mutations are classified as *passengers* because of their low rate of recurrence in clinical cancer samples that imply that they provide no proliferative benefit to cancer. Accumulated passengers have been previously assumed to be neutral^2,3^.

However, our recent analyses of human cancer genomics data ^4,5^ indicate that accumulated passengers, an inextricable consequence of genomic instability, can be moderately deleterious to cancer cells possibly because they alter important functional protein sites. Furthermore, evolutionary modeling, where individual cells can acquire advantageous drivers and deleterious (to cancer cells) passengers, finds that these deleterious passengers largely evade natural selection (Figure 1). Although passengers exhibit individually weak effects on progression, their cumulative effect can commensurate with that of drivers because of their disproportionately high numbers. As such, passengers can reverse and prevent tumor growth in models where the population size of genetically unstable cells freely fluctuate (Figure 1). The predictions of this evolutionary modeling, however, have yet to be tested experimentally.

**Figure 1:**
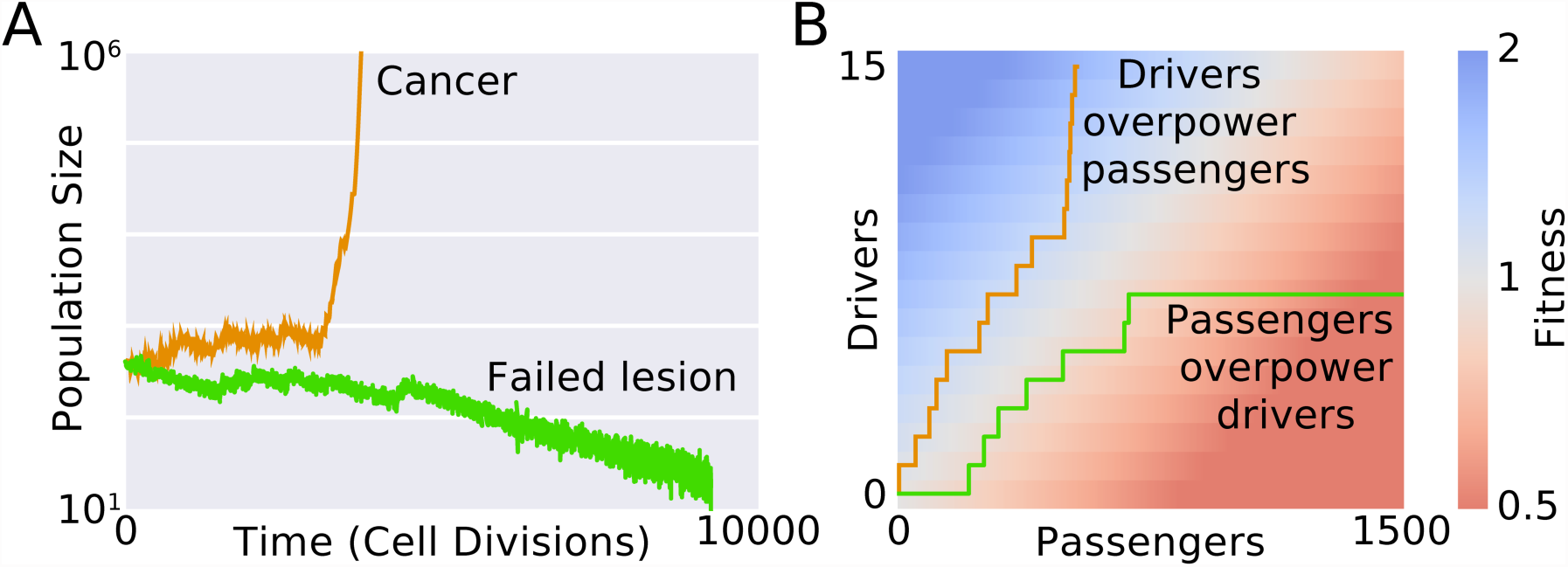
In our model, advantageous drivers compete with deleterious passenger. **A** Prior minimalistic modeling^1^ investigated cancer evolution where cells can mutate durring division, aquring either a strong advantageous drivers or (more frequently) a moderately-deleterious (to cancer) passengers. Population size freely fluctuates about a carrying capacity that is proportional to the evolving mean fitness of the population. In this model, initially identical trajectories diverge in population size, either progressing to cancer or regressing to extinction. **B** Tumor fitness and population size in this model is determined by the relative abundences of drivers and passengers. Successful tumors acquired drivers disproportionatly faster than passengers.

Other recent studies find that passengers, can increase tumor immunogenicity ^6–9^ and that very high genomic instability correlates with improved clinical outcomes^6,10^. In these immunogenic models, neoepitope passengers activate MHC-based immunoediting of tumors and immune checkpoint inhibitors. However, these studies focus on single nucleotide variants (ignoring the effects of chromosomal alterations) and do not directly test the effect of passengers on individual cell fitness, nor in a driver-controlled setting. Earlier studies^10,11^ also reported better clinical outcomes for patients with high rates of genomic instability. While providing a link between mutational load and clinical response, such studies were correlative and did not establish a causal effect of the passenger load on cancer.

Here we developed human cell line and mouse models to directly assess the effects of passenger load on cell fitness and carcinogenesis. We find that passengers are deleterious to cell fitness and tumor progression. We develop a precise metric of aggregate passenger load of chromosomal alterations, finding that passengers’ fitness cost can counterbalance and overcome cancer drivers in tumor populations. In mouse models, we demonstrate that genomic instability considerably slows tumor growth and that elevated passenger loads reduce metastatic burden. Our findings refute traditional paradigms of passengers in cancer as neutral events and suggest possible therapeutic avenues to exploit this vulnerability.

## Results

To directly test for the effects of passenger load on proliferative fitness in cancer cells, we developed cell lines with identical drivers and controlled passenger loads (Fig. 2A). The distribution and dispersion of accumulated passengers mimicked observed passengers in human cancer (Fig. 2E).

**Figure 2:**
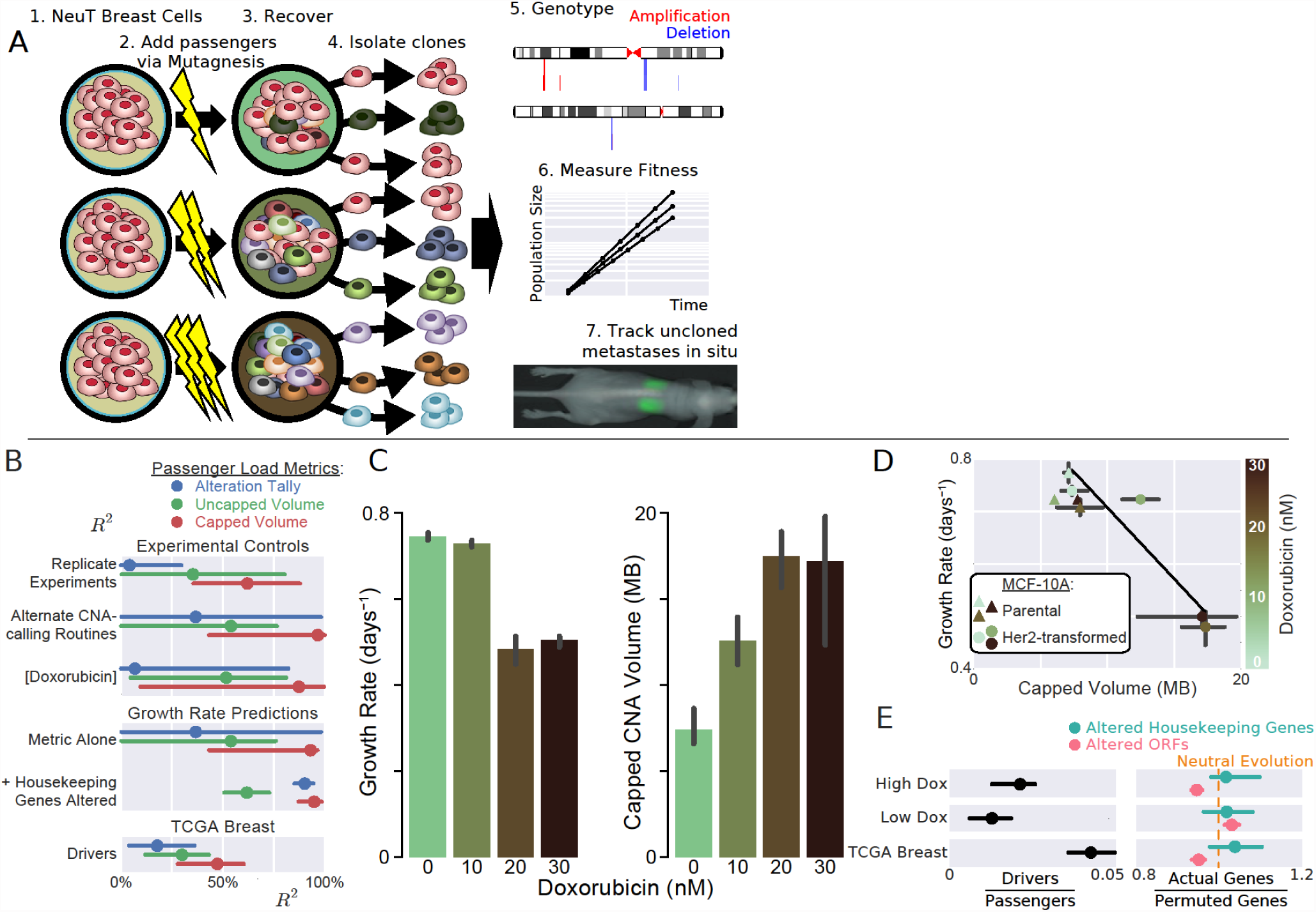
Passenger alterations reduce cell proliferation. **A** Increasing loads of passengers were introduced into Her2-transformed MCF-10A breast cells by mutagenic Doxorubicin. After a 2-week recovery, clones were then isolated, genotyped for Copy-Number Alterations (CNAs), and assayed for cell fitness and metastatic potential. **B** Three measures of passenger load (a CNA Tally, Uncapped Volume, and Capped Volume) were studied for their ability to predict various experimental controls, cell line fitness effects, and properties of TCGA genomes. Capped Volume proved to be the most consistent between (i) replicate experiments and (ii) the parameters of our CNA-identification algorithm^2^ (SI Methods), while also (iii) correlating best with Doxorubicin exposure. It was also most negatively correlated with growth rate. We considered a simple linear model between passenger metric and growth rate, as well as a multi-linear model that included the quantity of altered housekeeping genes in each sample as a second predictor. Lastly, our evolutionary modeling predicts that passenger load will linearly correlated with the number of drivers in sequenced human cancers, a phenomenon that was most pronounced in Capped CNA Volume^3^. **C** Increasing dosages of mutagen decreases proliferative potential and increases passenger load. **D** The fitness cost of passengers in transformed cells was -0.028 MB^-1^ (r^2^ = 84%, 95% CI: 64—99%). Untransformed cells neither acquired passengers nor decreased in proliferative potential, suggesting that passenger accumulation, and not doxorubicin toxicity, causes fitness reduction. **E** (Left) Passenger alterations dominated the genomes of our mutagenized cell lines, as intended (Low Dox = 0/10 nM Doxorubicin; High Dox = 20/30 nM Doxorubicin). (Right) Passenger alterations spanned housekeeping genes and Open Reading Frames (ORFs) *nearly* as often as expected by random chance in both our cell lines and genuine human breast cancers. If natural selection prevented deleterious passengers from accumulating, then a depletion of CNAs spanning particularly important genomic regions would be expected. The neutral rate (random probability) of CNAs spanning a gene set was determined by randomly permuting the locations of each gene in the set 200 times. Error bars (95% CI) and p-values throughout this study were calculated using bias-corrected bootstrapping^4^.

First, spontaneously-immortalized, and genomically-stable human breast epithelial cells (MCF-10A) were transformed with a single driver—activated Her2/Erbb2 oncogene—using a retroviral expression system^12,13^. To ensure genomic homogeneity of the initial population, an individual clone was isolated after transformation and alterations in this clone were subtracted from subsequent genomic analysis (Fig. S1). Then, cells from this clone were treated with different doses of doxorubicin at sub-lethal, sub-clinical levels (0 – 30 nM) overnight. Doxorubicin, being an inhibitor of TOPO2, introduces copy-number alterations, mimicking natural genomic instability^14^. Cell lines were then given a 2-week recovery period to eliminate any residual doxorubicin toxicity. Individual clones were isolated from each mutagenized population to ensure genomic homogeneity within each sample (Fig. 2A). As a control, un-transformed MCF10A cells were subjected to the same protocol.

Genotyping of the developed cell lines, using a combination of SNP-array and low-coverage whole-genome sequencing (Fig. S1), confirmed that (i) increasing doxorubicin levels incur greater quantities of passenger alterations and that (ii) very few additional drivers accumulated (mean 1.4/sample, Fig. 2E& Table S1). Her2-transformed clones acquired on average 296 novel passenger alterations following treatment with 20 or 30nM doxorubicin—significantly higher than the parental untransformed MCF10A cells (p < 0.005, Table S1). Driver alterations, which were rare and did not vary substantially between samples, were identified as either amplifications spanning a known breast cancer oncogene, or deletions spanning a known tumor suppressor (Methods, Table S2). Thus, these clones allowed us to explore fitness landscapes with nearly-identical and rare drivers and increasing passenger loads.

In agreement with our hypothesis, increasing passenger loads negatively correlated with doubling time; cell lines with the highest passenger load (20 nM and 30 nM Doxorubicin exposure) grew >30% slower than un-mutagenized strains (Fig. 2C). Doubling times were measured by daily cell counting on plates for three days. Doxorubicin exposure did not alter the growth of untransformed MCF-10A cells (with functioning DNA repair) and accumulated few additional passengers, demonstrating that mutagen exposure alone cannot reduce growth and that passengers are required (Fig. 2D).

Taken together these results show that proliferative fitness of cancer cells declines with the load of accumulated passengers.

To test whether the patterns of accumulated passenger alterations in our study were consistent with genuine passenger alterations observed in cancer, we analyzed cancer genomics data. Negative selection against passengers was minute in both our model and genotyped breast cancers (Fig. 2E). The probability that a passenger altered a housekeeping gene was similar between our samples, clinical breast cancers, and randomly-permuted data. Thus, the passengers in this study are representative of clinical passengers and do not include mutations that would not accumulate in cancer.

Next we asked whether passenger load could quantitatively predict fitness costs. Because a simple tally of passengers ignores their dramatic variation in length and ploidy^15,16^, we considered two alternative measures: *CNA Volume,* which weights Copy-Number Alterations by their length and change in ploidy: 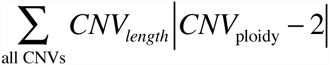; and *Capped CNA Volume,* which caps individual CAN lengths in the summation above at 2 MB. Conceptually, CNA volume denotes the total (genome-wide) deviation from normal gene dosage. We capped CNAs lengths because their length distribution dramatically varies with their origin: *focal* CNAs, which begin and end within a chromosomal arm, are categorically shorter than *non-focal* CNAs, which contain a centrosomal or telomeric terminus^15,16^ (Fig. S2A). Non-focal CNAs dramatically sway CNA Volume if uncapped and may not sway fitness by such a profound amount, perhaps because centromeres and telomeres contain fewer functional elements. Capped CNA Volume outperformed Uncapped CNA Volume (which outperformed a simple tally) in its ability to robustly summarize passenger load by being most consistent between (i) replicate experiments and (ii) alteration-calling algorithms, and (iii) most correlated with mutagenic dose (Fig. 2B& S2B). Not surprisingly, growth rates also correlated most negatively with this metric (r^2^ = 84%, 95% CI: 64–99%); however, all metrics negatively correlated with growth rate (p < 0.05, Figure S3), so metric choice does not engender a particular result.

We further validated these characteristics of passenger load using clinical breast cancer data. Earlier we predicted and observed a positive linear relationship between passenger load and the number of drivers^5^ that results from deleterious passengers in cancer genomes being counterbalanced by additional drivers to yield equivalent tumor fitness. We found that our introduced measures of passenger load, CNA Volume (and *Capped* CNA volume in particular), exhibited stronger linear relationships with driver events (Fig. 2B). Collectively, these results further establish cancer development as a tug-of-war between drivers and passenger, and Capped CNA Volume as a superior measure of passenger load.

Developed cell lines that carry differential passenger loads and the same drivers allow us to directly measure the fitness cost of passenger alterations. By regressing the growth rate to the Capped CNA Volume we find the mean fitness cost of -0.027 per MB, 95% CI [-0.0213, -0.056] (Figure 2E) i.e. ∼5% growth reduction per CNA longer than 2 Mb, and proportional to length with ∼3% per Mb for focal (<2Mb) CNAs. This measurement is in excellent agreement with our earlier inference of ∼3% of fitness loss per Mb for CNA in human cancers (2%-10% depending on the chromosome), which was based on analysis of ∼40,000 intra-chromosomal-arm CNAs from more than 3,000 cancer specimens and Chromosome Conformation Capture, Hi-C, data^15^.

Measured CNA fitness cost allows us to estimate the total passenger load of human breast cancers. We calculate that an average breast cancer contains a Capped CNA Volume of 146 MB, suggesting that passengers dramatically hinder tumor growth–by >300%. While seemingly high, both our previous evolutionary modeling and genomic analyses are consistent with passengers having such a high impact on cancer growth. Modeling predicts that drivers and passengers are in a delicate balance, where the total fitness cost of passengers is barely outweighed by the collective effect of drivers. To test this, we analyzed human breast cancer genomes and found a strong linear relationship between drivers and passengers (Figure S2C). Indeed, tumors with an average passenger load contain 151% more drivers than an extrapolated tumor without any passengers. Cancers may also evolve mechanisms to compensate for this extreme passenger load^17^; in silico modeling of tissue hierarchies finds that chromosome losses are ∼5 times less costly in cancer cells than normal tissue^18^. Nonetheless, our prior evolutionary simulations demonstrate that passengers in this range of fitness costs often accumulate, can overpower drivers, and offer an effective therapeutic target (see Discussion)^4^. Taken together results of the cell line experiment combined with genomic analysis show that cancers carry a very substantial passenger load that can significantly reduce their fitness.

To investigate passenger fitness effects in a broader context, emphasizing clinical utility, we turned to mouse cancer models. We first used transgenic mice to investigate the effects of increased genomic instability (inducible by cytotoxic^19^ and targeted^20^ chemotherapies) on tumor growth. Unlike traditional paradigms where genomic instability always accelerates carcinogenesis^1,21^, we predicted that tumor growth can be slowed or even suppressed when mutation rates exceed a critical level^5^ (SI). To test this prediction, we created mouse models of breast carcinogenesis with high and low mutation rates by crossing a MMTVneu mouse model of Her2-positive breast cancer^13^ (mice containing a single driver— activated Her2 (NeuT) expressed in the mammary epithelium), with mice containing a homozygous deletion of histone H2AX that is necessary for DNA double-strand break repair^12^ (SI). Hybrid progeny carried a single copy of the NeuT oncogene and were H2AX haploinsufficient. As a control, we used animals that also carried a single copy of NeuT and both copies of H2AX gene.

Tumors emerged after a median of 10 months (Figure 3A). We then isolated and genotyped 7 tumors using low-coverage sequencing (Methods), which affirmed that tumors in H2AX^+/-^ mice accumulated passengers at a higher rate than in control mice (p = 0.11, Figure 3C). Tumors with elevated mutation rates (H2AX^+/-^) emerge at approximately the same time (possibly somewhat later, *p* = 0.05) and grew significantly slower (*p* < 0.001) than control tumors (Figure 3), thereby demonstrating that (i) passengers are deleterious in organismal environments, and (ii) genomic instability can be detrimental to tumor growth ^1,22^.

**Figure 3:**
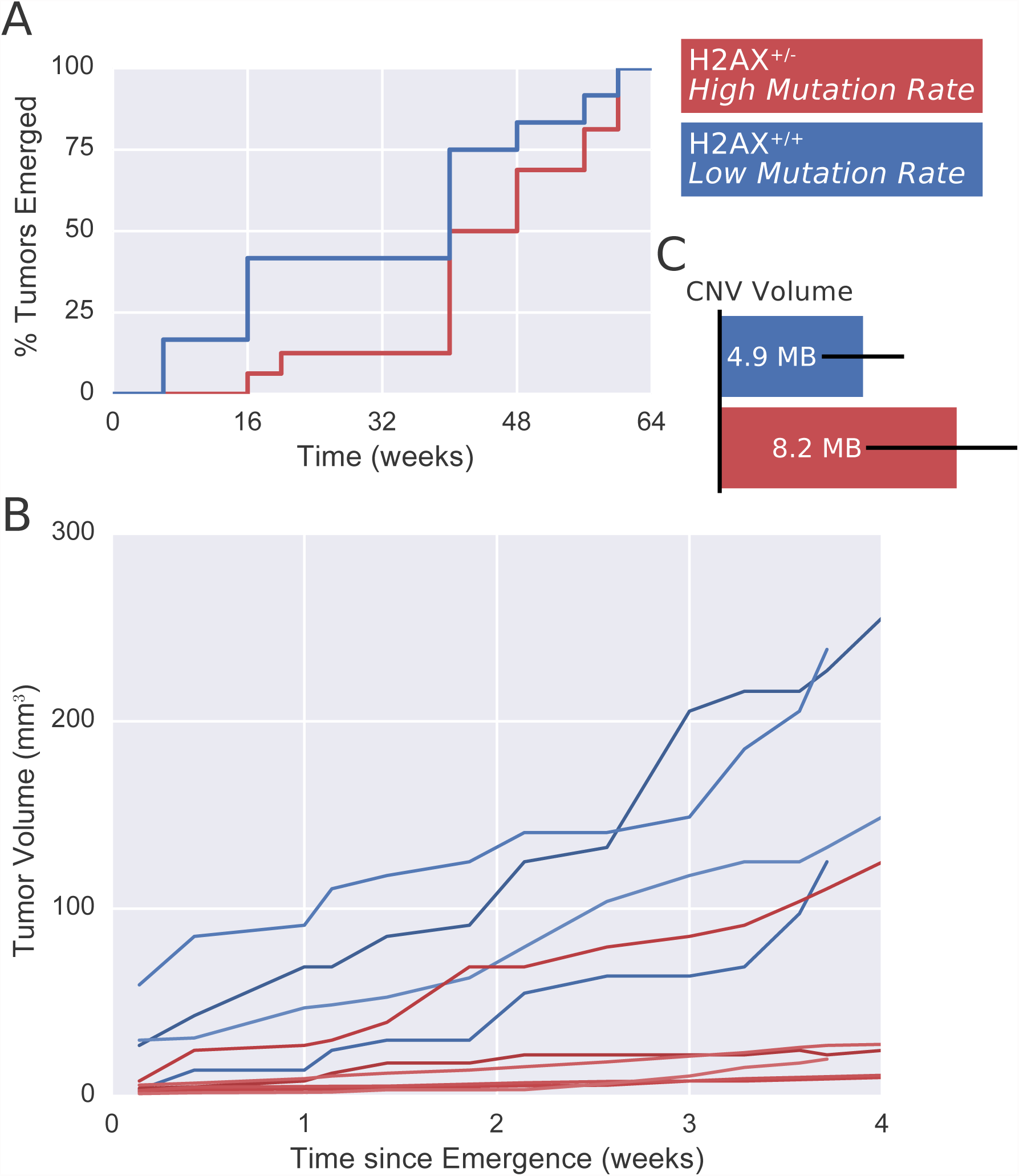
Elevated mutation rates slow tumor growth in a mouse model. **A** Breast cancers, instigated by Her2-transformation, emerged from mice in highly-mutagenic (H2AX^+/-^) conditions at similar times as moderate-mutagenic (H2AX^+/+^) conditions (possibly later, *p* = 0.05). **B** Tumors with elevated mutation rates grew 52% slower (95% CI: 40—66%) after emergence. **C** Genotyping confirmed that elevated mutations rates increase passenger loads.

Our model makes additional predictions regarding the tug-of-war between drivers and passengers that we then tested. Because the H2AX^+/-^ tumors grew slower, their mutation rate should exceed the critical rate and, thus also exceed clinical breast cancer mutation rates (SI). Indeed, the H2AX^+/-^ tumors mean mutation rate (28 MB/y 95% CI: 13—44) was ∼10x greater than human breast cancers (2.7 MB/y 95% CI: 2.3—3.2). Also, the passenger:driver ratio should increase in H2AX^+/-^ tumors^5^, which indeed was the case (Table S2). Drivers, in this analysis, were amplified/deleted mouse homologs to the human oncogenes/tumor suppressors identified above (Table S3). This analysis provides direct support for a tug-of-war between drivers and passengers in an organismal environment during cancer development.

Next we asked whether passengers impacted not only primary tumor growth, but also metastatic development. Our recent simulations indicate that deleterious passengers interfere with metastatic evolution since additional passengers accumulate during the population bottleneck, which occurs during dissemination and micrometastatic evolution of individual cells. Because our transgenic mouse model does not metastasize, it could not be used to assess effects of passengers on metastasis. We, therefore, investigated passengers’ effect on metastatic progression by (i) introducing a luciferase reporter into developed Her2-transformed MCF10A cells, with increasing passenger loads (see above), (ii) injecting these cells into the tail vein of SCID female mice, and (iii) monitoring lung metastases in situ via thoracic bioluminescence. Metastases arose after week 4 (Figure 4A) in all groups. After 7 weeks, the number of metastases in the mutagenized (10/20 nM Doxorubicin exposure) strains was 12-fold lower (95% CI: 9–19) then in the control group, while aggregate metastatic load (bioluminescence) was 80-fold (95% CI: 54–128) lower (Figure 4BC). These results indicate that passenger load dramatically impacts metastasis, reducing both the efficiency of new metastasis formation and their progression.

**Figure 4:**
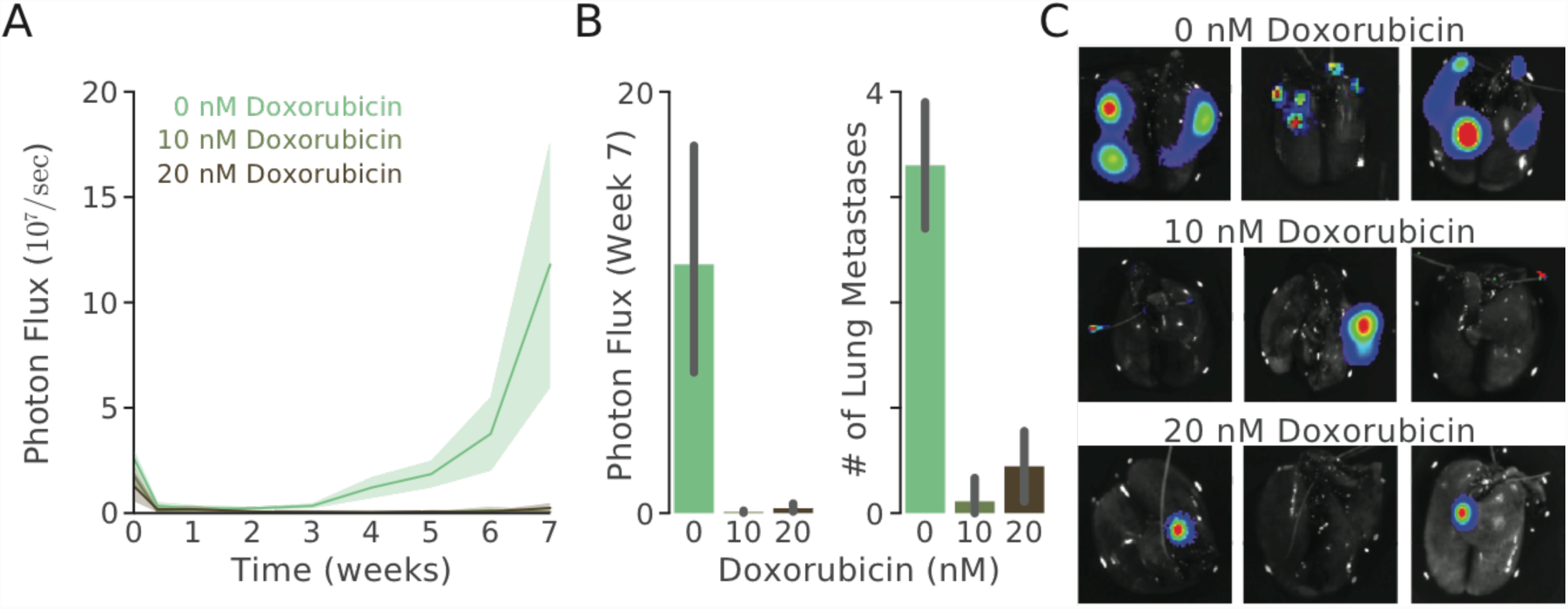
**A**Her2-transformed, MCF-10A breast cells were mutagenized as before, transfected with a Luciferase reporter, and injected into the tail-vein of SCID mice (10 per line) without cloning. Lung metastases, were monitored in situ by live cell bioluminescence imaging for 7 weeks. **B** At 7-weeks, total metastatic burden and the number of lung metastases (identified by dissection and counting) were dramatically greater in unmutagenized cells. **C** Representative ex-vivo bioluminescent lung images.

## Discussion

Our findings demonstrate that accumulated passengers can be directly deleterious to cancer, by reducing cell proliferative fitness, significantly slowing down cancer growth, and impeding the formation and growth of metastases. Overall, our findings support several unique predictions of evolutionary models incorporating deleterious passengers (Table 1) and bolster the prior modeling and human-genomic evidences of deleterious passengers in cancer. In this new paradigm, genomic instability is a double-edged sword: it accelerates driver events, but eventually accumulates intolerable quantities of deleterious passengers. Indeed, if genomic instability had no disadvantage, then evolving drug resistance would become impossibly easy. Defeating cancer requires understanding, and corralling, its tempo and mode of evolution.

**Table 1.** Predictions of the Deleterious Passenger Model supported by these experiments. Our experiments investigated four key predictions of the Deleterious Passengers Model that were previously established using evolutionary simulations, theoretical analysis, and genomic evidences (detailed in the SI).

| Predictions of the<br>Deleterious Passenger Model |
| --- |
| ~0.1-1% fitness cost per passenger |
| Aggregate passenger load can overpower<br>drivers—preventing tumor progression |
| Exceptionally high mutation rates inhibits<br>carcinogenesis |
| Deleterious passengers especially limit<br>metastatic progression |

Deleterious passengers are clinically relevant as a diagnostic and as a targetable phenotype. Prior results indicated possible targetable immunogenic effects of passengers ^6–9^ that can supplement their direct effects. We showed that passenger load directly affects overall tumor fitness (in the absence of immunoediting), including its metastatic capacity. Segregating passenger (i.e. subclonal passengers) may also interfere with the acquisition of adaptive drivers^23^ and reduce genetic diversity^24,25^, thereby preventing drug resistance. Hence, deleterious passengers may affect both tumor’s current aggressiveness and future evolution.

Accurately characterizing passenger load is essential to any clinical effort that exploits passengers. Passenger load is an ideal genomic biomarkers because its large quantity mitigates error in assessing individual mutation’s fitness costs. Nonetheless, we improved assessments by weighting passenger’s impact by their length and ploidy. Genetic context mattered less: the number of altered genes (or, specifically, housekeeping genes) correlated only moderately with fitness reduction, as seen previously ^15^, and only minutely improved our ability to predict fitness reduction in our cell lines (p = 0.04, Fig. 2B).

Therapeutics can target cancer’s deleterious passenger load by increasing the passenger mutation rate (discussed above) or by increasing passenger deleteriousness. Passengers are deleterious because they may (i) disrupt essential cell functions^26^; (ii) induce cytotoxic stress via protein misfolding, dis-balance, and aggregation^11^; or (iii) create immune-eliciting neoantigens^6^. All three of these phenotypes can be exacerbated and, as modeling shows, lead to cancer meltdown. Our experimental findings clinically-benefit these efforts by (a) identifying a biometric to use in directing such therapies, and (b) suggesting that these treatments will work best in metastatic prevention and in conjunction with mutagenic therapies. We hope to soon begin testing these hypotheses in the clinic.

